# Inhibition of methylthioadenosine phosphorylase provides protection from experimental acute kidney injury

**DOI:** 10.1101/2025.02.19.639065

**Authors:** Afaf Saliba, Yidong Chen, Jonathan W. Nelson, Abhinav Vetcha, Wei Wei Wang, Li Kang, Nagarjunachary Ragi, Soumya Maity, Hamid Rabb, W. Brian Reeves, Kumar Sharma

**Affiliations:** Center for Precision Medicine, University of Texas Health Science Center at San Antonio; San Antonio, TX 78229, USA; Division of Nephrology, Department of Medicine, University of Texas Health Science Center at San Antonio; San Antonio, TX 78229, USA; Department of Population Health Sciences, University of Texas Health Science Center at San Antonio; San Antonio, TX 78229, USA; Greehey Children’s Cancer Research Institute, University of Texas Health Science Center at San Antonio; San Antonio, TX 78229, USA; Division of Nephrology and Hypertension, Department of Medicine, Keck School of Medicine of University of Southern California. Los Angeles, CA 90033, USA; Department of Medicine, Johns Hopkins University, Ross 970, 720 Rutland Avenue, Baltimore, MD, 21205, USA

**Author notes:** **CORRESPONDING AUTHOR** Kumar Sharma, MD, Address: 7703 Floyd curl Drive, San Antonio, Texas 78229, USA.

## Abstract

Acute kidney injury (AKI) increases mortality risk and predisposes individuals to chronic kidney disease. Metabolic pathways play a crucial role in AKI pathophysiology. Here, we investigate the potential of methylthioadenosine phosphorylase (MTAP) inhibition as a novel renoprotective strategy in AKI. Using AKI mouse models, we demonstrate that a small molecule MTAP inhibitor significantly reduces kidney injury markers and improves renal histology. RNA sequencing reveals that MTAP inhibition modulates pathways associated with inflammation, oxidative phosphorylation, and cell survival. Additionally, analysis of human single-cell RNA sequencing data links MTAP expression to kidney injury marker in AKI. This study provides evidence of MTAP inhibition as a potential therapeutic strategy for AKI, highlighting metabolic dysregulation as a target for future clinical interventions.

## INTRODUCTION

Acute kidney injury (AKI) in native kidneys occurs in 2-5% of inpatients, increases mortality, and can lead to chronic kidney disease (CKD) ^1^. Experimental studies have demonstrated that agents targeting metabolic pathways and p53 have therapeutic potential ^2,3^. Recent clinical studies have suggested that sodium-glucose linked transport inhibitors (SGLT2i) could have benefit for reducing the incidence of AKI ^4^. These findings underline the importance of metabolic pathways in AKI and suggest opportunities to explore additional therapeutic strategies targeting cellular stress responses and metabolic dysregulation, regardless of diabetic status.

We recently demonstrated a renal protective effect of methylthioadenosine phosphorylase (MTAP) inhibition in experimental models of diabetic kidney disease ^5^. MTAP is an enzyme involved in purine and polyamine metabolism, breaking methylthioadenosine (MTA) down for nucleotide recycling. Here, we explored effects of MTAP inhibition in AKI mouse models. We found that a small molecule MTAP inhibitor (MTAPi) significantly reduced markers of kidney injury and improved renal histology. RNA sequencing (RNA-seq) identified pathways critical for kidney health, such as inflammation, oxidative phosphorylation, and cell survival. Analysis of human single-cell RNA-seq (scRNA-seq) data correlated MTAP expression to kidney injury marker expression in AKI. This is the first study to identify the benefit of MTAP inhibition in AKI, laying the foundation for future research on mechanisms and clinical translation.

## METHODS

Male 10-12-week-old C57BL/6J mice were used to model ischemic and cisplatin-induced AKI. MT-DADMe-ImmA, a small molecule MTAPi was administered intraperitoneally. Kidney injury was assessed by renal biomarkers, histology and RNA-seq. ScRNA-seq data from 32 human kidney biopsies (20 healthy, 12 AKI) retrieved from Kidney Precision Medicine Project (KPMP) ^6^ were analyzed. Detailed methods are provided in the **Supplementary Material**.

## RESULTS

### MTAP inhibition protects against ischemic and cisplatin-induced AKI

MTAPi or vehicle control was administered to mice prior to renal ischemia-reperfusion (IR). MTAPi treatment mitigated AKI, as demonstrated by markedly improved BUN (**Figure 1A**). The expression of hepatitis A virus cellular receptor 1 (*Havcr1*) and lipocalin-2 (*Lcn2*) was significantly attenuated in the MTAPi-treated group (**Figure 1, B-C**). Periodic acid-Schiff (PAS) staining of kidney tissues showed a significant decrease in the number of necrotic tubules in the MTAPi-treated-IR mice (**Figure 1D)**. MTA level significantly increased in the plasma and kidney cortex of MTAPi-treated-IR mice. MTAP metabolizes MTA to produce adenine and 5-methylthioribose-1-phosphate the decreased adenine-to-MTA ratio and MTA accumulation indicate MTAP inhibition (**Supplementary Figure S1**).

**Figure 1.**
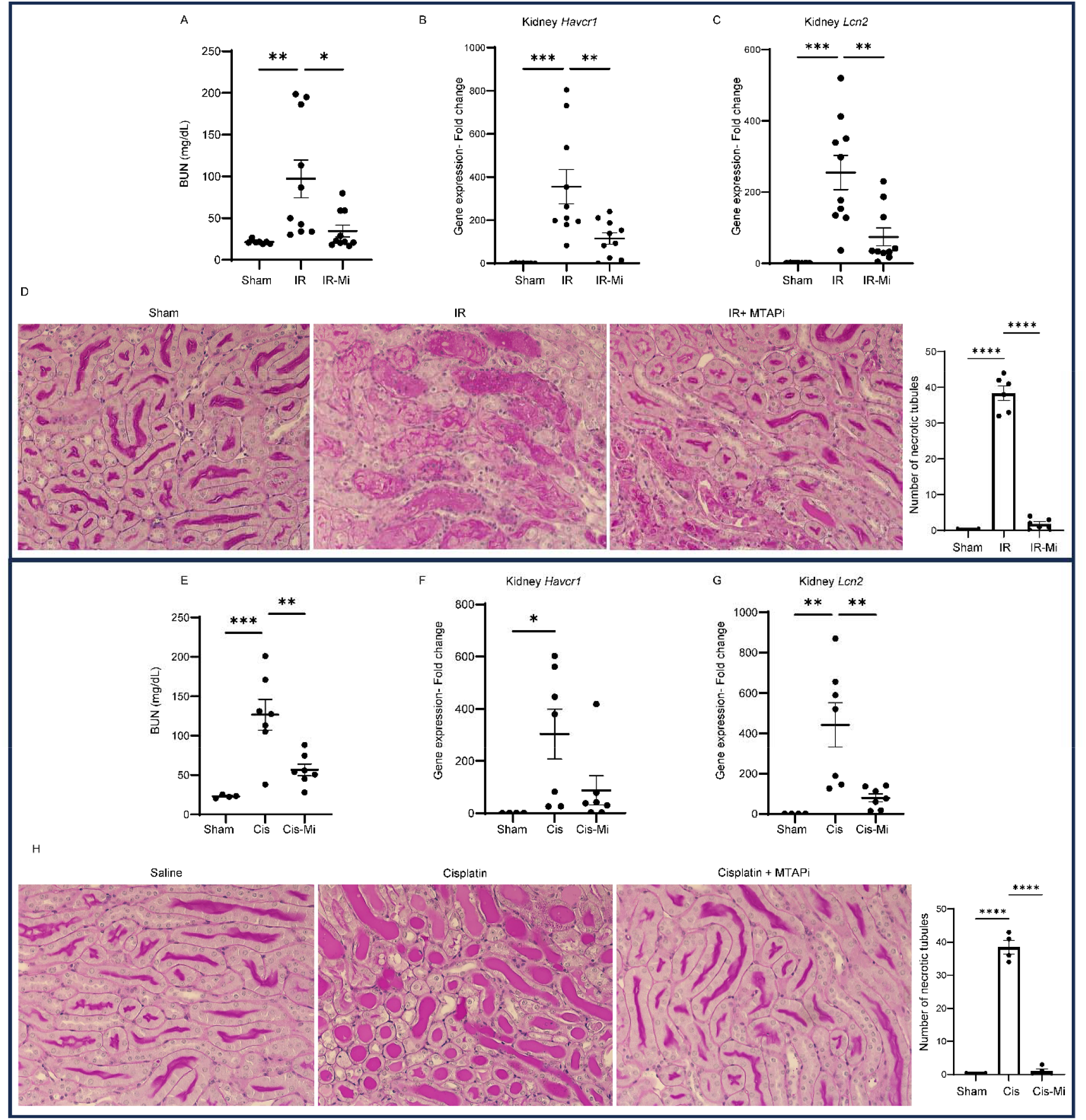
MTAP inhibition mitigates acute kidney injury in ischemic and cisplatin AKI models. Male mice were treated with MTAPi (20 mg/kg) or vehicle (n=10 per group) before bilateral IR, sham-operated mice (n=7) as controls. **(A)** Blood urea nitrogen (BUN) levels and qPCR analysis of kidney cortex RNA lysates **(B)** mRNA levels of hepatitis A virus cellular receptor 1 (*Havcr1)* (known as Kim1) and **(C)** Lipocalin 2 (*Lcn2)* (known as NGAL), **(D)** PAS staining of kidney sections from sham (n=4), IR-vehicle (n=6), and IR-MTAPi (n=6) with corresponding quantification of necrotic tubules per average of 20 fields (40x). Male mice were administered MTAPi (20 mg/kg) or vehicle (n=7 per group) 4.5h prior and 24, and 48h post-receiving cisplatin (20 mg/kg) IP injection. Control mice received saline (n=4). **(E)** BUN levels, **(F)** mRNA levels of *Havcr1*, **(G)** *Lcn2*, and **(H)** PAS staining of kidney FFPE sections (n=4 per group) were evaluated with corresponding quantification of necrotic tubules per average of 20 fields (40x). Gene fold qPCR analyses were normalized to *Gapdh* levels. Graphs display means +/-SEM; *: p < 0.05, **: p < 0.01, ***: p < 0.001, ****: p < 0.0001, determined with one-way ANOVA followed by Tukey multiple comparison tests.

To assess whether MTAPi could be protective against other forms of AKI, we examined its effects in a cisplatin-induced AKI mouse model. MTAPi administered before and during cisplatin administration mitigated kidney injury, as evidenced by significant reduction of BUN, *Havcr1*, and *Lcn2* levels (**Figure 1, E-G**). MTAPi reduced tubular necrosis in histological analysis (**Figure 1H**).

### MTAP inhibition reduced inflammatory and stress response pathways in AKI

To gain mechanistic insight into MTAPi protection during AKI, bulk RNA-seq analysis of kidney cortex lysates revealed 2,063 differentially expressed genes (DEGs) in IR-vehicle *vs*. sham, while MTAPi pre-treatment reduced DEGs to 247 (**Supplementary Figure S2, A and Supplementary Table S2**).

Principal component analysis (PCA) showed distinct clustering, with IR-MTAPi samples closer to sham than IR-vehicle. DEG analysis confirmed this pattern (**Supplementary Figure S2**). MTAPi significantly downregulated inflammatory pathways, including TNFα/NFκB signaling, IL6-JAK-STAT3 signaling, TGF-β signaling, and p53 signaling, while upregulating fatty acid metabolism and oxidative phosphorylation when compared to IR-vehicle. Additionally, MYC target pathways, which were strongly upregulated during ischemia, were suppressed by MTAPi. (**Figure 2 A and Supplementary Figure S3**). Gene ontology (GO) biological process analysis showed MTAPi rescued pathways like stress response, cell death regulation, and metabolic processes (*e*.*g*., organic acid, small molecule, oxoacid, carboxylic acid, and amino acid metabolism) (**Figure 2 B and Supplementary Figure S4**). GO molecular function analysis further highlighted preserved transporter activity, oxidoreductase function, and adhesion molecule interactions. Cellular component analysis showed that MTAPi restored epithelial integrity-related gene expression, enriching apical/basolateral plasma membrane, brush border, mitochondria and peroxisome-associated genes which were disrupted in IR *vs*. sham.

**Figure 2.**
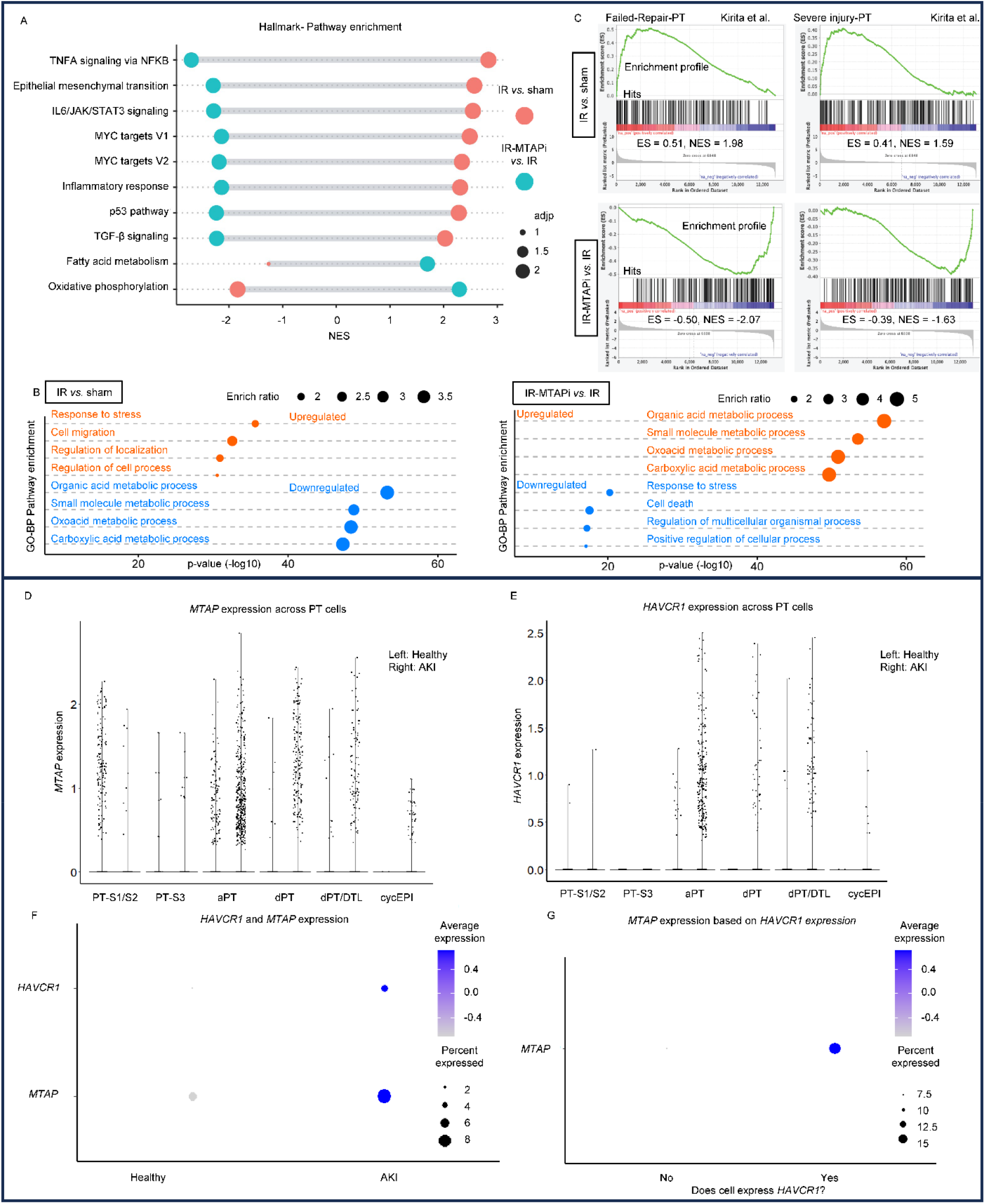
Comprehensive RNA-seq data analysis reveals that MTAP inhibition in mice reduces factors associated with proximal tubule injury and that MTAP is abundant in HAVCR1 positive proximal tubules in humans with AKI. **(A)** Dumpbell plot displaying the normalized enrichment scores (NES) for top pathways (Hallmark). Blue: IR-vehicle *vs*. sham and pink: IR-MTAPi *vs*. IR-vehicle **(B)** Dot chart of the top pathways *via* gene ontology (GO) biological process (BP) pathway enrichment analysis. Left: IR-vehicle *vs*. sham and right: IR-MTAPi *vs*. IR-vehicle. (C) GSEA enrichment plots show the correlation of differentially expressed genes in IR-vehicle *vs*. Sham (Top) and IR-MTAPi vs. IR (bottom) with failed repair PT (left) and severely injured PT (right) (data set from Kirita et al. ^7^). NES, Normalized Enrichment Score; ES, Enrichment Score. **(D)** *MTAP* expression across PT subpopulations under Healthy (Left) *vs*. AKI conditions (Right) **(E)** *HAVCR1* expression across PT subpopulations under Healthy (Left) *vs*. AKI conditions (Right) **(F)** Multi-dimensional dot plot representation of *HAVCR1* and *MTAP* expression in healthy *vs*. AKI samples. **(G)** Multi-dimensional dot plot representation of *MTAP* expression based on *HAVCR1*. The analysis shown in D-G was based on scRNA-seq data from the KPMP subset PT subpopulations comparing Healthy and AKI patients’ samples. PT-S1/S2: Proximal tubule cells from the S1 and S2 segments; PT-S3: Proximal tubule cells from the S3 segment; aPT: adaptive PT cells; dPT: dedifferentiated proximal tubule cells; and dPT/DTL: dedifferentiated PT/descending thin limb cells; cycEPI: cycling epithelial cells.

Concurrently, there was a suppression of extracellular-matrix-related gene expression (**Supplementary Figure S5 and S6**). Interestingly, while genes altered by IR were positively correlating with the failed repair and severe injured proximal tubules (PTs) genes signature ^7^ MTAPi showed a significant negative correlation (**Figure 2C)**. Overall, kidney bulk RNA-seq data suggested a reduction in stress-induced metabolic reprogramming and maladaptive repair, processes that are often triggered in kidney injury and specifically in PTs during injury ^7,8^.

### *MTAP* abundance in *HAVCR1*-expressing cells highlights its potential role during AKI

To determine the relevance of MTAP in the clinical manifestations of human kidney disease we investigated the scRNA-seq dataset from the KPMP ^6^. Because the PT is a key site of injury after AKI ^7^, we focused our analysis of the KPMP dataset for this segment of the nephron wherein the PT cells were annotated for both functional and pathological PT cell-states (**Supplementary Figure S7, A-B**). Within PT cells, *MTAP* expression was highest in adaptive PT (aPT) cells and moderately expressed in degenerative PT (dPT) (**Figure 2D**), and more abundantly expressed in samples from patients with AKI, indicative of injury-related transcriptional changes. aPT cells exhibit stress-response markers and are linked to successful and failed tubular repair ^6^. *MTAP* expression was significantly elevated in AKI samples compared to healthy references, remaining enriched in PT cells but also in DTL, ATL/TAL and PEC populations, suggesting a role for MTAP in response to injury (**Supplementary Figure S7, C**). Consistent with what is previously known about AKI ^7^, *HAVCR1*, an established marker of kidney injury, exhibited increased expression in samples from AKI patients, which paralleled MTAP expression (**Figure 2E**). In AKI, both *HAVCR1* and *MTAP* significantly increased compared to healthy reference (**Figure 2F**). To further establish a link between kidney injury and MTAP, we classified cells based on the expression of *HAVCR1* and found that *MTAP* was more abundant in *HAVCR1*-expressing cells (**Figure 2G**). Collectively, these findings underscore the role of MTAP in mediating the transcriptional landscape of PT cells and their broader involvement in the renal injury response during AKI.

## DISCUSSION

We demonstrate that targeting MTAP with a small-molecule inhibitor protects against both ischemic and nephrotoxic AKI in mice. Blocking MTAP resulted in significant MTA accumulation at the plasma and kidney levels which may provide renal protection possibly by inducing vasodilation *via* adenosine A2B receptor activation ^*9*^. Analysis of human scRNA seq data revealed an association between MTAP and HAVCR1 expression in human kidneys, with HAVCR1-positive PTs in AKI exhibiting high *MTAP* levels, suggesting its involvement in acute stress responses. While the precise mechanisms remain unclear, we hypothesize that AKI triggers MTAP activity, exacerbating cellular stress and damage.

MTAP was found significantly upregulated in aPTs which Lake et al. have identified as a distinct injury-response population with characteristic expression of *HAVCR1* and that correlate with failed tubular repair in rodents ^6^. Kirita et al. highlighted subsets of PT cells in murine AKI kidneys with severe injury and failed repair characteristics as metabolically dysregulated, shifting to inflammatory and profibrotic transcriptional programs^7^, a pattern mirrored in our IR dataset and prevented by MTAPi. Additionally, Liu et al. identified several PT-specific genes (Aspg, Pm20d1, Bhmt2, Slc7a13, Thnsl2) downregulated in IR^8^, many of which we found were preserved by MTAPi (**Supplementary Table S3 and S4**), reinforcing its role in targeting PT dysfunction. Together, these findings highlight a previously unrecognized role for MTAP in worsening kidney injury and position MTAP inhibition as a promising therapeutic strategy to mitigate PT dysfunction in AKI.

## ACKNOWLEDGMENTS

We thank the team of the Center for Precision Medicine for peer-review, data discussions, and technical support with mass spectrometry. Data was generated in the Genome Sequencing Facility which is supported by the University of Texas Health Science Center at San Antonio, NIH-NCI P30 CA054174 (Cancer Center at UT Health San Antonio) and NIH Shared Instrument grant S10OD030311 (S10 grant to NovaSeq 6000 System), and CPRIT Core Facility Award (RP220662). This work was supported by the National Institutes of Health, National Center for Advancing Translational Sciences grant TL1 TR002647 to AS. National Institutes of Health, National Heart, Lung, and Blood Institute grant T32 HL007446 (AS). University of Texas Health Science Center at San Antonio, Long School of Medicine-Research, Multi-PI pilot grant 2023-2024 (KS). National Institutes of Health grant UO1DK114920 (KS) Veteran’s Affairs Merit grant I01BX001340 (KS) and DoD CDMRP grant PR181598 (KS). Results in Figure 2 are in part based upon data generated by the Kidney Precision Medicine Project. Accessed November 20th, 2024. https://www.kpmp.org.

## AUTHOR CONTRIBUTIONS

Conceptualization and methodology: AS and KS; Investigation: AS with technical help and support from AV, WWW, LK and NR; Formal Analysis and visualization: AS, YC and JN; Resources: WBR, KS; Funding acquisition: AS, KS; Supervision: KS; Writing – original draft: AS; Writing – review & editing: AS, JN, SM, HR, WBR, KS.

## DATA SHARING STATEMENT

RNA-seq data are under preparation for submission to NCBI/GEO for data repository for public release. All other data are available in the main text or the supplementary materials. Bioinformatic code for KPMP datasets analysis and figure generation has been deposited on GitHub (https://github.com/JWNelsonLab).

