## Supplementary materials for "Inhibition of methylthioadenosine phosphorylase provides protection from experimental acute kidney injury"

Saliba et al.

**Supplementary Materials**

**Methods**

**Supplementary Figures**

Supplementary Figure S1. Impact of methylthioadenosine phosphorylase (MTAP) inhibition on methylthioadenosine (MTA) levels.

Supplementary Figure S2. Comparative analysis of kidney cortex RNA-seq data in ischemic-AKI model.

Supplementary Figure S3. Gene set enrichment analysis (GSEA).

Supplementary Figure S4. Gene ontology (GO) biological process (BP) enrichment analysis.

Supplementary Figure S5. Gene ontology (GO) molecular function (MF) enrichment analysis.

Supplementary Figure S6. Gene ontology (GO) cellular component (CC) enrichment analysis.

Supplementary Figure S7: Single-cell RNA-sequencing analysis of human kidney in AKI.

**Supplementary Tables**

Supplementary Table S1. Primers list for qPCR

Supplementary Table S2. Number of differentially expressed genes (DEG) between groups

Supplementary Table S3. Top DEGs IR30-Vehicle *vs.* Sham

Supplementary Table S4. Top DEGs IR30-MTAPi *vs.* IR30-Vehicle

**Supplementary References**

### Methods

Ischemic AKI models

All animal studies were IACUC-approved at the University of Texas Health Science Center at San Antonio. Ten- to twelve-week-old male C57BL/6J mice (The Jackson laboratory; #000664) were used for the study. Mice were anesthetized by inhalation of isoflurane (3% in oxygen). All procedures were performed on anesthetized mice on controlled heating pads with body temperatures monitored and maintained at 36-37 °C. Bilateral renal ischemia-reperfusion (IR) was conducted as previously described [1] with some modifications. Briefly, the renal pedicles were identified *via* flank incisions and clamped for 30 min with small nontraumatic vascular aneurysm clips (Roboz Surgical Instruments, Gaithersburg, MD). After clamp removal, reperfusion was confirmed by observing restoration of kidney color, indicating arterial blood flow. Sham-operated mice underwent an identical procedure with no pedicle clamping. Incisions were closed in two layers. 0.5 ml warm saline was given subcutaneously after the procedure to compensate for the loss of body fluid during surgery. Mice were euthanized at 24h after ischemic injury.

Cisplatin-induced AKI model

C57BL/6J male mice (8-12 weeks old) (The Jackson laboratory; #000664) were injected intraperitoneally (IP) with 20 mg/kg cisplatin dissolved in saline solution, as described previously[2,3]. Mice were euthanized 72h later, and plasma and organs were collected.

Treatment protocols and criteria

The MTAP inhibitor (MTAPi), MT-DADMe-ImmA (AdooQ BioScience; # A13651), was dissolved in 10% DMSO in 1XPBS, and the vehicle was prepared using 10% DMSO in 1XPBS. The MTAPi (20 mg/kg) or vehicle was administered by intraperitoneal injection. The time and dosage of administration were selected to achieve optimal inhibition of MTAP activity based on previous studies [4], the t1/2 for the onset of inhibition was 50 min, with complete inhibition achieved by 250 min. To ensure sustained inhibition of MTAP during the renal ischemia-reperfusion procedure, we administered the MTAPi intraperitoneally 4.5h before the induction of ischemia or before cisplatin injection. Cisplatin treated mice received an additional dose of MTAPi or vehicle 24 and 48h post-cisplatin injection.

Histology

Kidneys were paraffin-embedded, and 5-micron sections were stained with PAS. The necrotic tubule score per section represents the average of scores noted from 20 randomly selected regions within the section. Counting of necrotic tubules were done in a blinded manner by experienced research scientists.

Liquid chromatography/mass spectrometry (LC-MS)

Selective metabolites were measured from plasma, or frozen kidney cortex lysates by LC-MS. LC-MS analyses were performed on the Thermo Q Exactive HF-X Orbitrap mass spectrometer (Thermo, San Jose, CA) interfaced with heated electrospray ionization source (HESI) and coupled with Thermo Vanquish HPLC system. An aliquot of 2.5 µL of the sample was injected into the instrument using an autosampler. The chromatographic separation took place on an Agilent ZORBAX HILIC PLUS column with 3.5µm particle size and with the dimensions 2.1 ×100 mm with a phase composition of 10 mM ammonium formate, 0.05% formic acid in Millipore water (component A), and 0.05% formic acid in 95% acetonitrile (component B) at 0.3 mL/ min flow rate was used [5].

RNA sequencing (RNA-seq) and bioinformatics analysis

Total RNA was extracted from the homogenized kidney cortex samples (6x sham, 6x IR, 6x IR, and 5x IR-MTAPi) using the QIAGEN RNeasy Mini Kit (Cat# 74104) according to the manufacturer's instructions RNA samples were further processed for RNA-seq library at Greehey Children’s Cancer Research Institute (GCCRI) Genome Sequencing Facility (GSF) following the NEB Directional RNA library preparation guide with NEBNext rRNA depletion kit v2 (human/mouse/rat) (New England Biolabs, Ipswich, MA). The first step in the workflow involved the depletion of rRNA by hybridization of complementary DNA oligonucleotides, followed by treatment with RNase H and DNase to remove rRNA duplexed to DNA and original DNA oligonucleotides, respectively. Following rRNA removal, the RNA was purified, fragmented, and copied into first strand cDNA using reverse transcriptase and random primers, followed by the 2nd strand cDNA synthesis using DNA Polymerase I and RNase H. The cDNA fragments then went through an end repair process followed by ligation of adapters. The products were then purified and enriched by PCR amplification to generate final RNA-seq libraries, which were sequenced using Illumina NovaSeq 6000 platform at 100 bp paired-end module at GCCRI GSF. Sequence reads (average of ~29 million paired reads per sample) were aligned to USCS mm10 mouse genome using HISAT2 aligner (v2.1.0)[6] followed with Stringtie (v2.1.3b) to quantify gene expression levels (both in read counts and in Fragment Per Kilobase of transcript (FPKM) for all NCBI RefSeq genes[6]. Differential gene expression analyses were performed using DESeq (R/Bioconductor) for each group *vs*. sham group, as well as between IR and IR-Mi [7]. Differentially Expressed Genes (DEGs) were determined by the following criteria: (1) absolute log2 fold change >1, (2) one of the average FPKM of comparison groups >1, and (3) multiple-test adjusted P value < 0.05 (Benjamini-Hochberg method). We also performed Gene Set Enrichment Analysis (GSEA,[8] ) by using fgsea implementation in R/Bioconductor [9] using differential expression fold-change). Enriched functions were selected with p-value < 0.01 and false-discovery rates (FDR). Genes with selected functions were further plotted in heatmaps using R/pheatmap function (genes with very low expression were removed).

Quantitative polymerase chain reaction (qPCR)

Total RNAs from homogenized tissue samples were extracted using the QIAGEN RNeasy Mini Kit (Cat# 74104) according to the manufacturer's instructions. cDNA synthesis was performed using the Thermo Fisher Scientific RevertAid Reverse Transcription Kit (Cat# 4374966). The primer sequences for the target genes are shown in (Supplementary Table S1). qPCR was performed using QuantStudio3 with SYBR Green PCR Master Mix kit (Thermo Fisher Scientific, Cat# A25742).

Analysis of KPMP Datasets

The single-cell RNAseq data1 (PREMIERE_Alldatasets_08132021.h5Seurat) was downloaded from the KPMP website (KPMP.org). The clinical data provided on the KPMP website (OpenAccessClinicalData.csv) was then integrated into the meta data of the single-cell RNAseq Seurat object. A total of 32 biopsies were analyzed from 20 “healthy reference” samples and 12 “acute kidney injury” (AKI) samples. Proximal tubule cells were subsetted based on `subclass.l1 = PT` and healthy reference cells were subsetted based on `Enrollment.Category = "Healthy Reference" or "AKI"`. Proximal tubule cells were categorized based on the expression of Kim1 (gene = HAVCR1) using the code `ifelse(GetAssayData(KPMP_subset, assay = "RNA", slot = "data")["HAVCR1", ] > 0, "yes", "no")`. Single-cell RNAseq data was visualized using the DimPlot(), FeaturePlot(), and DotPlot() functions of the package Seurat (v4.4)2 in R (v4.3.1). Bioinformatic code for analysis and figure generation has been deposited on GitHub (https://github.com/JWNelsonLab)

**Supplementary Figure S1. Impact of methylthioadenosine phosphorylase (MTAP) inhibition on methylthioadenosine (MTA) levels.** (A) Schematic representation of MTAP activity. MTAP converts MTA into adenine and 5-methylthioribose-1-phosphate (MTR-1-P). MTAP inhibition increases MTA levels [10,11] and adenine/MTA (Ade/MTA) ratio is indicative of MTAP activity. MTA concentration in the (B) plasma and (C) kidney. Adenine-to-MTA ration (Ade/MTA) in the (D) plasma and (E) kidney. Graphs display means +/- SEM; *: p<0.05 and **: p<0.01 determined by two-tailed t-test.

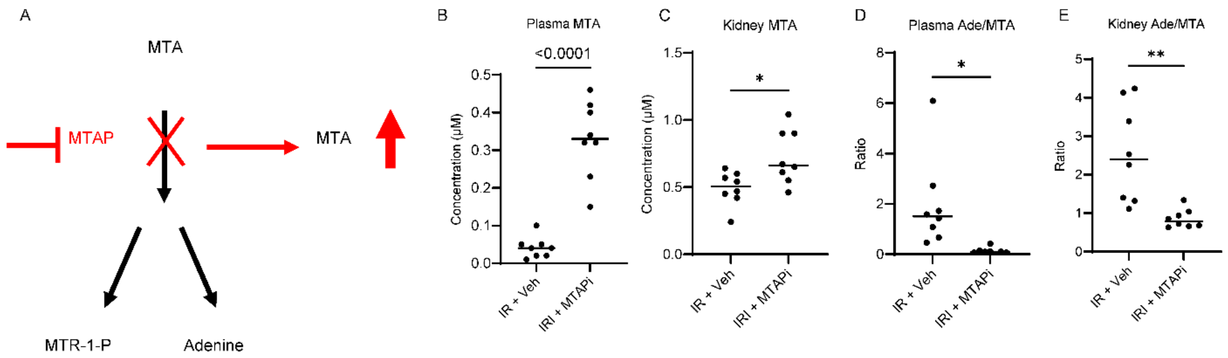

**Supplementary Figure S2. Comparative analysis of kidney cortex RNA-seq data in ischemic-AKI model.** RNA-seq with rRNA depletion was performed on kidney cortex RNA lysates. (A) Venn diagram displaying the overlap of differentially expressed genes (DEGs) among IR-vehicle *vs.* sham (green), IR-MTAPi *vs.* sham (pink), and IR-MTAPi *vs.* IR-vehicle (blue). (B) Principal Component Analysis (PCA) plot illustrating the distribution of samples based on principal components PC1 and PC2; IR-vehicle (yellow), IR-MTAPi (purple), and sham- (Blue). (C) Heatmap representation of DEGs across experimental conditions, illustrating the gene expression profiles of IR-vehicle (green), IR-MTAPi (orange), and sham groups (blue).

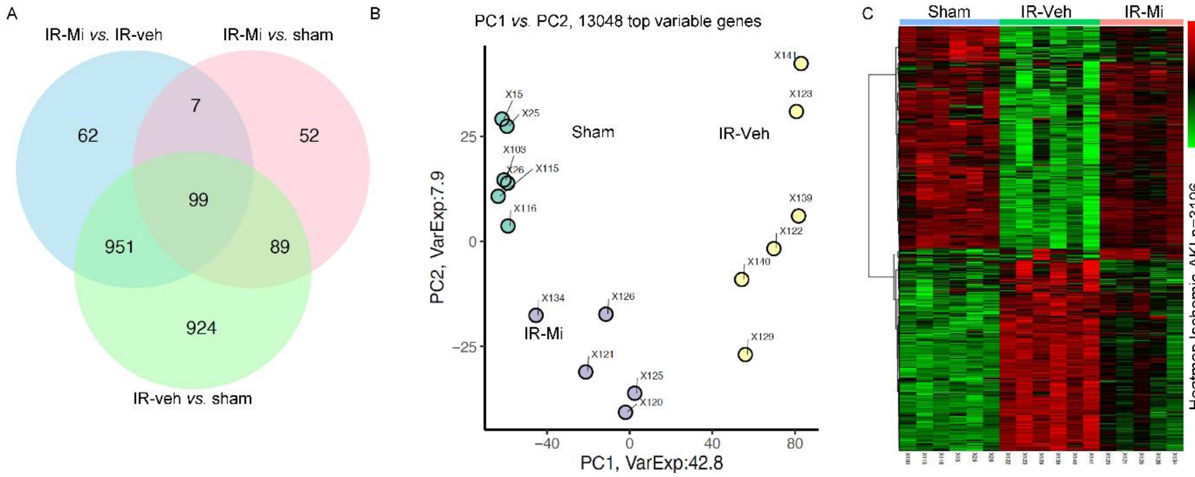

**Supplementary Figure S3. Gene set enrichment analysis (GSEA).** Top altered GSEA pathways (Hallmark database) in (**A**) IR-vehicle *vs*. sham and (**B**) IR-MTAPi *vs*. IR-vehicle. Related to Figure 2 A.

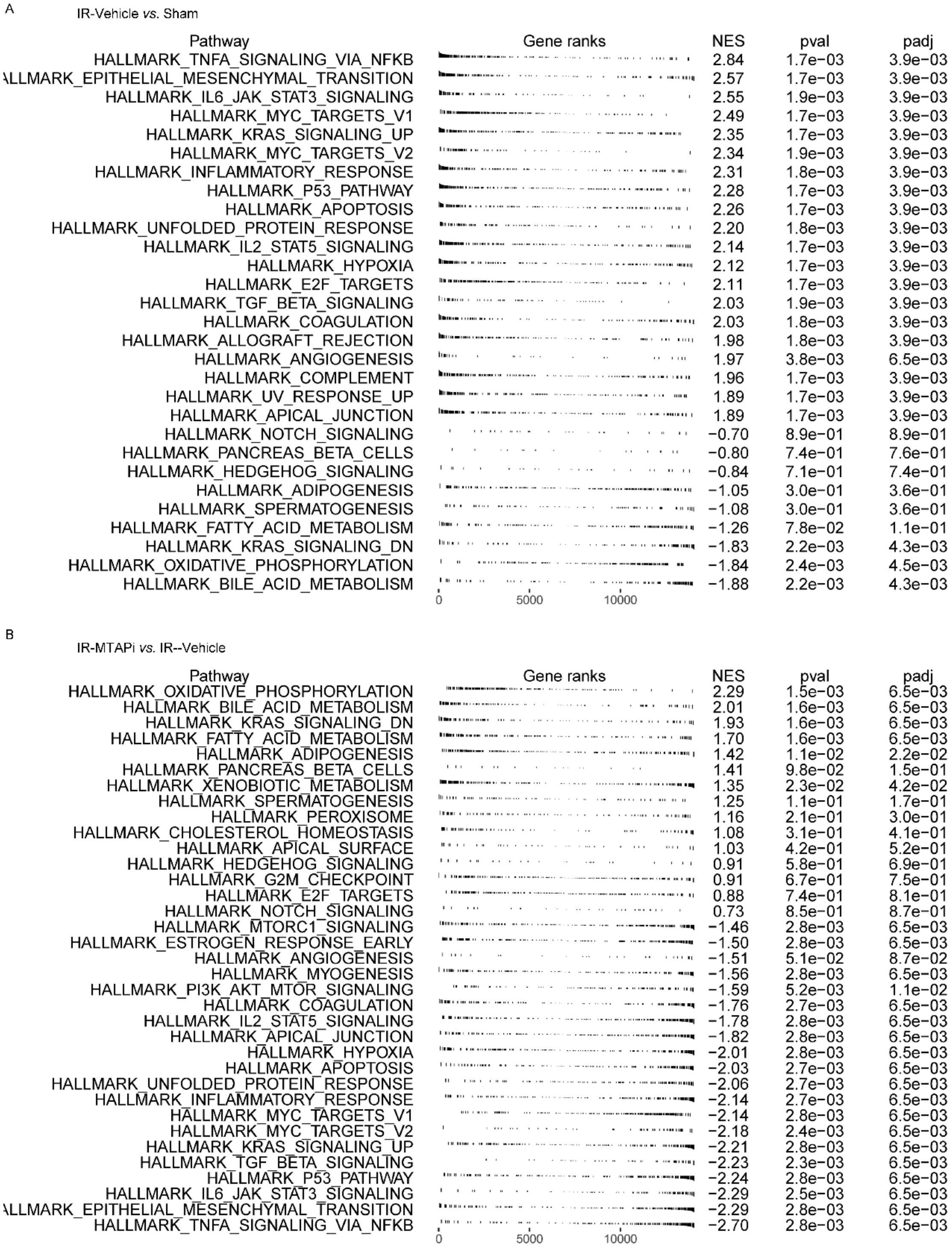

**Supplementary Figure S4. Gene ontology (GO) biological process (BP) enrichment analysis.** Dot chart of top 20 enriched biological processes identified *via* GO-BP analysis (**A**) IR-vehicle *vs.* sham-operated mice; (**B**) IR-MTAPi *vs.* IR-vehicle groups. Related to Figure 2 B.

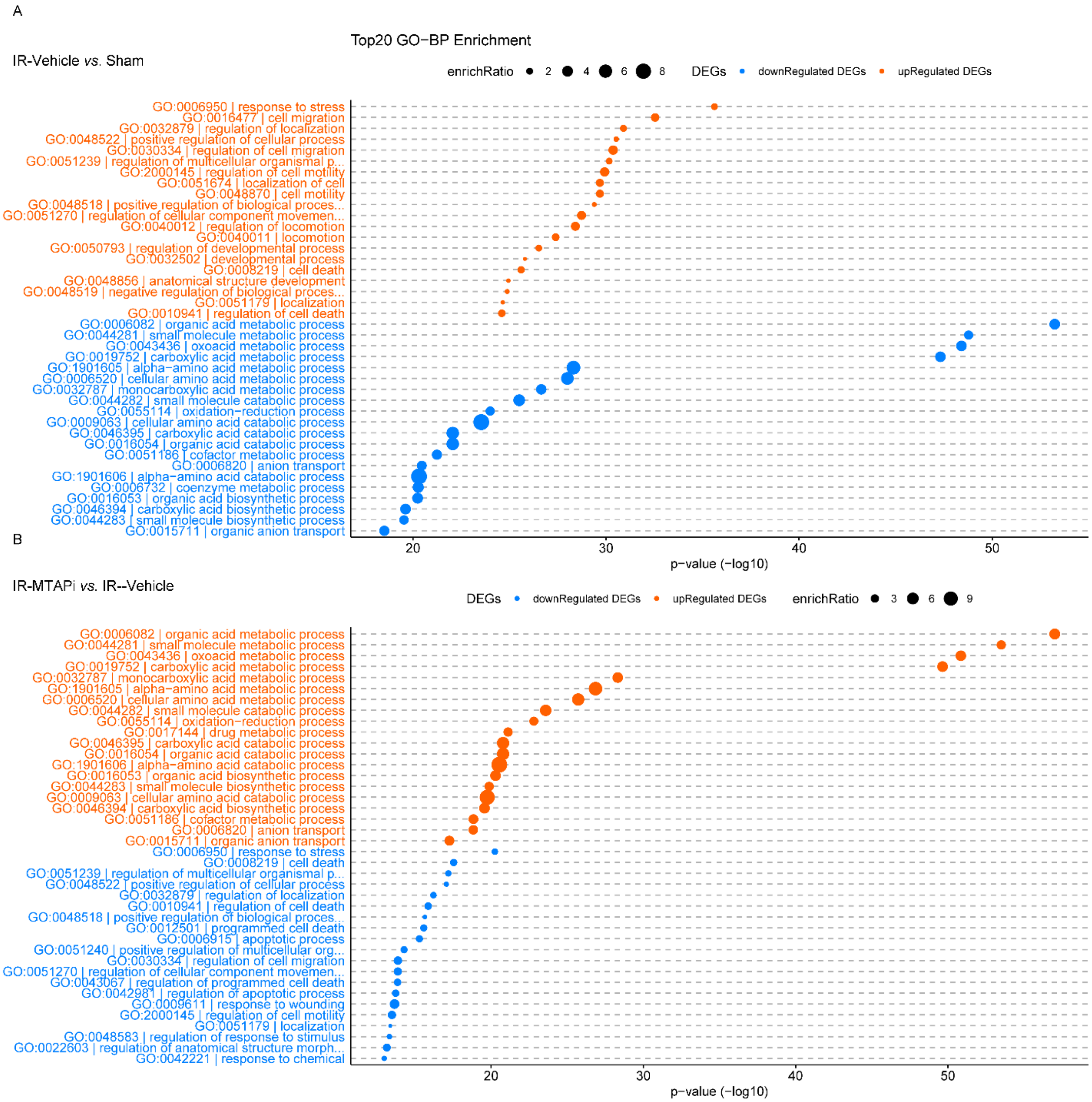

**Supplementary Figure S5. Gene ontology (GO) molecular function (MF) enrichment analysis.** Dot chart of the top 20 enriched molecular functions identified through GO-MF analysis (**A**) IR-vehicle *vs.* sham-operated mice; (**B**) IR-MTAPi *vs.* IR-vehicle groups.

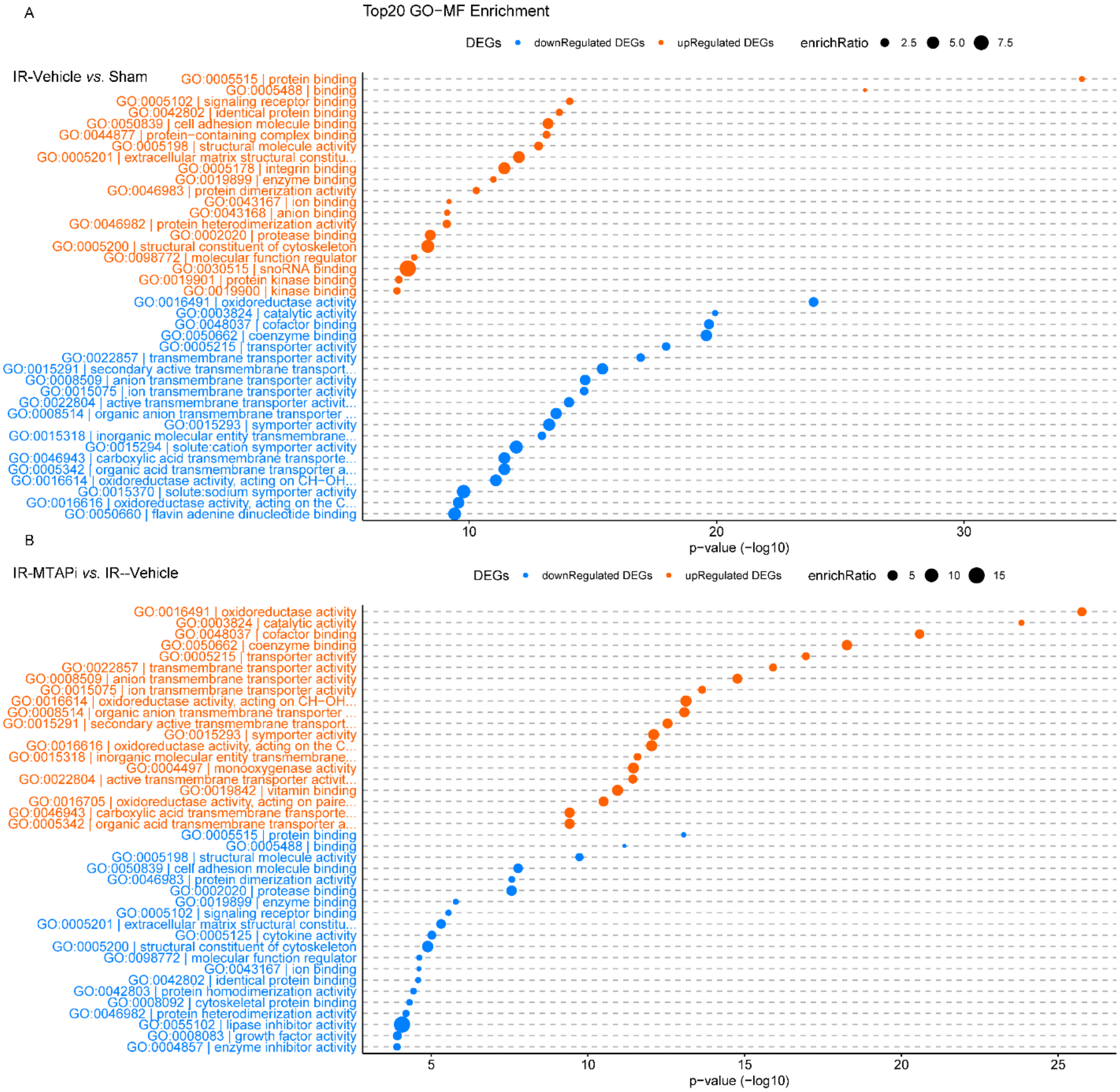

**Supplementary Figure S6. Gene ontology (GO) cellular component (CC) enrichment analysis.** Dot chart of the top 20 enriched cellular component processes identified through GO-CC analysis (**A**) IR-vehicle *vs.* sham-operated mice; (**B**) IR-MTAPi *vs.* IR-vehicle groups.

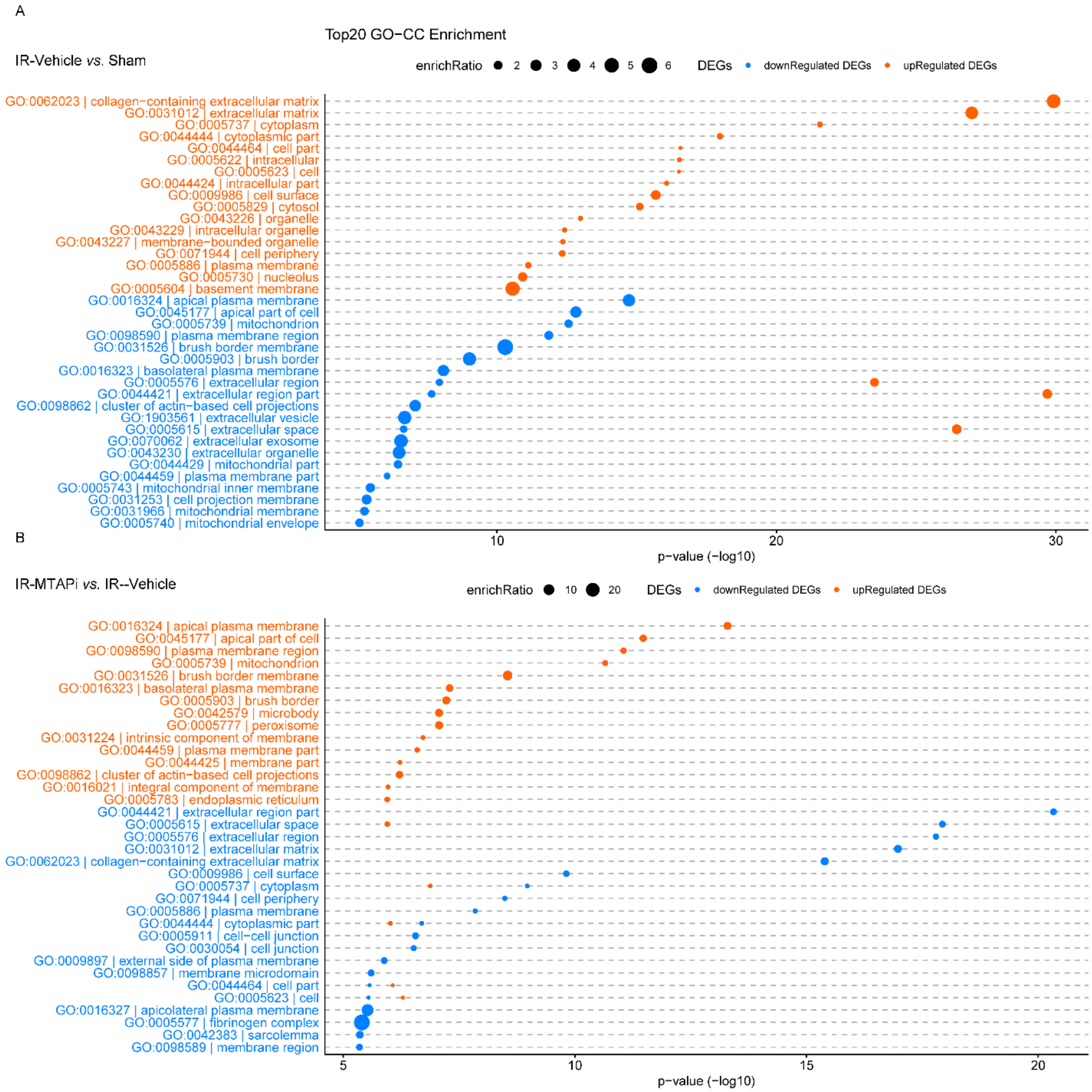

**Supplementary Figure S7: Single-cell RNA-sequencing analysis of human kidney in AKI.** (**A**) UMAP clustering of scRNA-seq data from the Kidney Precision Medicine Project (KPMP) (Lake B et al. 2023) [12]. We conducted additional analysis by isolating PT subpopulations in Healthy and AKI samples. UMAP plots (left) highlight these PT subsets. (**B**) UMAP plots showing PT cells clustering based on *HAVCR1* expression (top) and *HAVCR1* expression levels in PT cells (bottom). (**C**) Multi-dimensional dot plot of MTAP expression across kidney cell types in healthy (top) *vs.* AKI samples (bottom). PT: Proximal tubule, DTL: descending thin limb, ATL/TAL: ascending thin limb/thick ascending limb, DCT: distal convoluted tubule, CNT: connecting tubule, (PC): principal cells, IC: intercalated cells, POD: podocytes, PEC: parietal epithelial cells, EC: endothelial cells (EC).

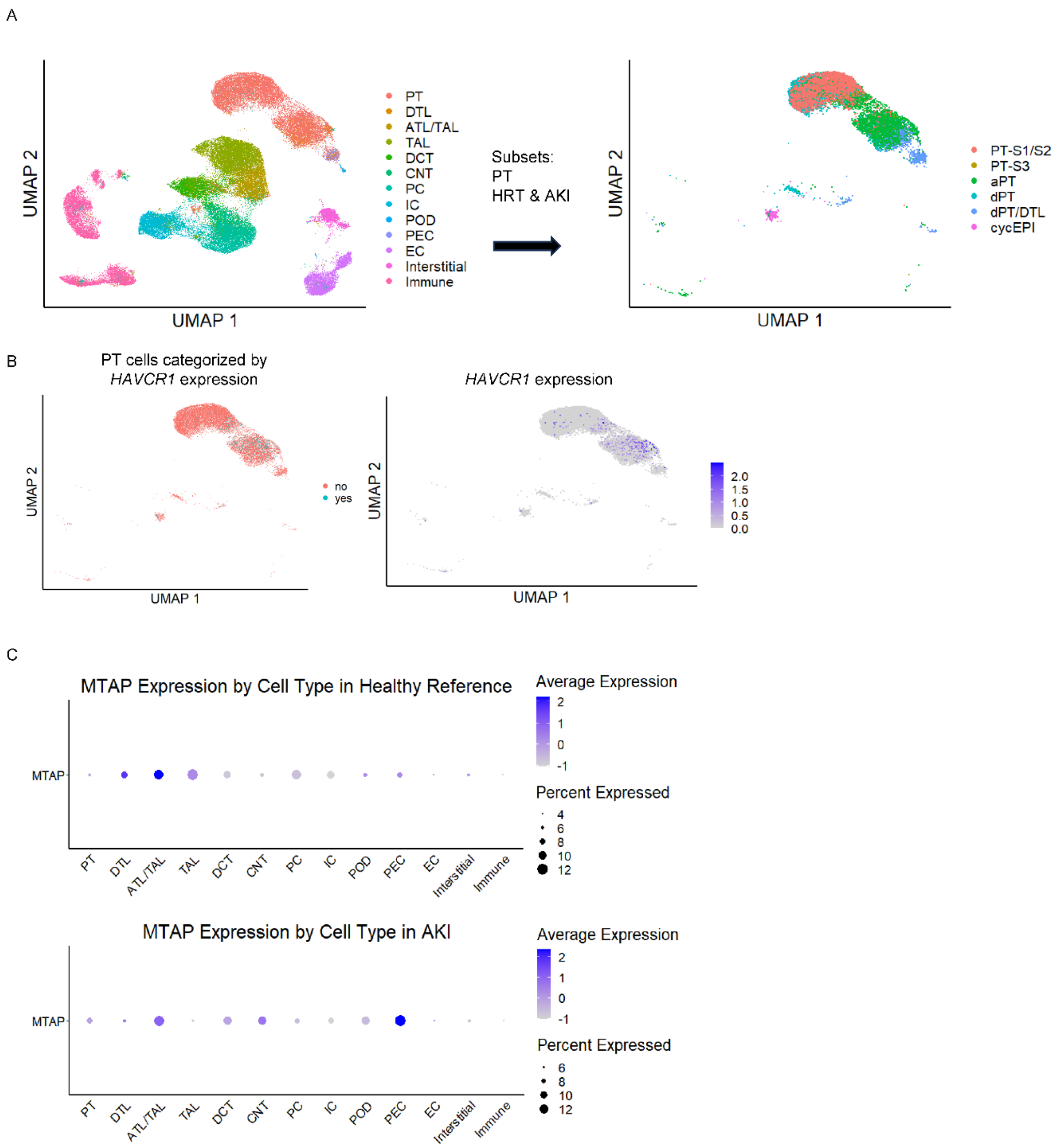


**Supplementary Figure S7: Single-cell RNA-sequencing analysis of human kidney in AKI.**

**Supplementary Table S1.** Primers list for qPCR

| Gene | Source | Forward | Reverse |
|---|---|---|---|
| Mouse-Gapdh | IDT | TGTGTCCGTCGTGGATCTGA | CCTGCTTCACCACCTTCTTGA |
| Mouse-*Havcr1* | IDT | CTCCAAGAAGACCCACAACTAC | GGAGGTAGAGACTCTGGTTGAT |
| Mouse-*Lcn2* | IDT | GCAGGTGGTACGTTGTGGG | CTCTTGTAGCTCATAGATGGTGC |

**Supplementary Table S2.** Number of differentially expressed genes (DEG) between groups

| Comparison | N (G1) | N (G2) | # of DEGs | Criterion |
|---|---|---|---|---|
| IR-vehicle *vs*. sham | 6 | 6 | 2063 | adj. p-value <= 0.05,<br>Fold-change >= 2.00 |
| IR-MTAPi *vs*. sham | 5 | 6 | 247 | |
| IR-MTAPi *vs*. IR-vehicle | 5 | 6 | 1119 | |

**Supplementary Table S3.** Top DEGs IR30-Vehicle *vs.* Sham

| Gene | log2FC | padj | |
|---|---|---|---|
| Havcr1 | 9.05 | 0.00e+00 | hepatitis A virus cellular receptor 1 |
| Hmox1 | 4.13 | 2.20e-111 | heme oxygenase 1 |
| Egf | -5.26 | 2.20e-111 | epidermal growth factor |
| Cml1 | -3.11 | 1.95e-110 | N-acetyltransferase 8 (GCN5-related) family member 1 |
| Ttc36 | -3.16 | 2.29e-106 | tetratricopeptide repeat domain 36 |
| Anxa2 | 3.39 | 3.72e-92 | annexin A2 |
| Atp4a | -3.21 | 6.72e-89 | ATPase, H+/K+ exchanging, gastric, alpha polypeptide |
| Smpdl3b | 4.69 | 1.98e-85 | sphingomyelin phosphodiesterase, acid-like 3B |
| Capg | 2.83 | 4.84e-84 | capping protein (actin filament), gelsolin-like |
| Akr1c21 | -2.96 | 8.20e-84 | aldo-keto reductase family 1, member C21 |
| Steap1 | 3.56 | 6.01e-83 | six transmembrane epithelial antigen of the prostate 1 |
| Lrp8 | 3.60 | 1.66e-79 | low density lipoprotein receptor-related protein 8 |
| Cyp2j11 | -2.57 | 2.92e-79 | cytochrome P450, family 2, subfamily j, polypeptide 11 |
| Myc | 3.62 | 7.58e-77 | myelocytomatosis oncogene |
| Slc16a7 | -2.50 | 2.76e-75 | solute carrier family 16 (monocarboxylic acid transporters), member 7 |
| Litaf | 2.47 | 3.23e-73 | LPS-induced TN factor |
| Aspg | -2.83 | 9.09e-73 | asparaginase |
| Pm20d1 | -2.51 | 9.13e-73 | peptidase M20 domain containing 1 |

**Supplementary Table S4**. Top DEGs IR30-MTAPi *vs.* IR30-Vehicle

| Gene | log2FC | padj | Gene name |
|---|---|---|---|
| Hsd3b3 | 2.60 | 1.05e-64 | hydroxy-delta-5-steroid dehydrogenase, 3 beta- and steroid delta-isomerase 3 |
| Pm20d1 | 2.13 | 5.69e-57 | peptidase M20 domain containing 1 |
| Havcr1 | -2.92 | 1.11e-45 | hepatitis A virus cellular receptor 1 |
| Bhmt2 | 1.81 | 1.60e-45 | betaine-homocysteine methyltransferase 2 |
| A4gnt | 3.68 | 2.41e-42 | alpha-1,4-N-acetylglucosaminyltransferase |
| Thnsl2 | 1.91 | 3.32e-42 | threonine synthase-like 2 (bacterial) |
| Hmox1 | -2.44 | 4.84e-41 | heme oxygenase 1 |
| Slc6a13 | 1.76 | 8.01e-40 | solute carrier family 6(neurotransmitter transporter,GABA),member 13 |
| Mep1b | 3.68 | 6.29e-38 | meprin 1 beta |
| Gyk | 1.88 | 1.65e-37 | glycerol kinase |
| Steap1 | -2.14 | 1.84e-37 | six transmembrane epithelial antigen of the prostate 1 |
| Aspg | 2.15 | 2.53e-37 | asparaginase |
| Slc16a9 | 1.61 | 2.21e-36 | solute carrier family 16(monocarboxylic acid transporters), member 9 |
| Spp2 | 1.59 | 1.69e-35 | secreted phosphoprotein 2 |
| Upb1 | 1.69 | 6.53e-35 | ureidopropionase, beta |
| Fads2 | 1.62 | 1.72e-34 | fatty acid desaturase 2 |
| Sectm1b | 2.68 | 7.63e-34 | secreted and transmembrane 1B |
| Abhd14b | 1.54 | 4.36e-33 | abhydrolase domain containing 14b |

10. Kirovski G, Stevens AP, Czech B, et al. Down-regulation of methylthioadenosine phosphorylase (MTAP) induces progression of hepatocellular carcinoma via accumulation of 5'-

deoxy-5'-methylthioadenosine (MTA). *Am J Pathol*. Mar 2011;178(3):1145-52. doi:10.1016/j.ajpath.2010.11.059

11. Sharma K, Zhang G, Hansen J, et al. Endogenous adenine mediates kidney injury in diabetic models and predicts diabetic kidney disease in patients. *J Clin Invest*. Oct 16 2023;133(20)doi:10.1172/JCI170341

12. Lake BB, Menon R, Winfree S, et al. An atlas of healthy and injured cell states and niches in the human kidney. *Nature*. Jul 2023;619(7970):585-594. doi:10.1038/s41586-023-05769-3
